# Aperiodic neural activity is a better predictor of schizophrenia than neural oscillations

**DOI:** 10.1101/113449

**Authors:** Erik J. Peterson, Burke Q. Rosen, Aysenil Belger, Bradley Voytek, Alana M. Campbell

**Affiliations:** Neurosciences Graduate Program, University of California, San Diego; 9500 Gilman Drive, La Jolla, 6 CA 92093, United States; Department of Cognitive Science, University of California, San Diego; 9500 Gilman Drive, La Jolla, 6 CA 92093, United States; Department of Psychiatry, University of North Carolina at Chapel Hill, Chapel Hill, NC 27599, United States; Department of Psychology, Carnegie Mellon University, Pittsburgh, PA 15213, United States

**Keywords:** schizophrenia, biomarker, aperiodic activity, classification

## Abstract

Diagnosis and symptom severity in schizophrenia are associated with irregularities across neural oscillatory frequency bands, including theta, alpha, beta, and gamma. However, electroencephalographic signals consist of both periodic and aperiodic activity characterized by the (1/f^X^) shape in the power spectrum. In this paper we investigated oscillatory and aperiodic activity differences between patients with schizophrenia and healthy controls during a target detection task. Separation into periodic and aperiodic components revealed that the steepness of the power spectrum better predicted group status than traditional band-limited oscillatory power in a classification analysis. Aperiodic activity also outperformed the predictions made using participants’ behavioral responses. Additionally, the differences in aperiodic activity were highly consistent across all electrodes. In sum, compared to oscillations the aperiodic activity appears to be a more accurate and more robust way to differentiate patients with schizophrenia from healthy controls.

**Significance statement:** Understanding the neurobiological origins of schizophrenia and identifying reliable and consistent biomarkers are of critical importance to improving treatment of that disease. Numerous studies have reported disruptions to neural oscillations in patients with schizophrenia. This has, in part, led to schizophrenia being characterized as a disease of disrupted neural coordination, reflected by changes in frequency band power. We report however that changes in the aperiodic signal can also predict clinical status. Unlike band-limited power though, aperiodic activity predicts status better than participants’ own behavioral performance and acts as a consistent predictor across all electrodes. Alterations in the aperiodic signal are consistent with well-established inhibitory neuron dysfunctions associated with schizophrenia, allowing for a direct link between noninvasive EEG and chronic, widespread, neurobiological deficits.

## Introduction

Schizophrenia is characterized by disruptions to multiple cognitive and behavioral domains, including significant working and long-term memory disruptions, impaired attention, and disorganized patterns of thought and language. These cognitive deficits are a special symptom domain, separate from the domains of delusions, hallucinations, and affective blunting^1,2^. Differences in single brain regions, or cognitive systems, cannot explain complex schizophrenic symptomatology. Instead, widespread physiological abnormalities--leading to reduced and disorganized neural communications--are thought to be a better explanation for and predictor of schizophrenic pathophysiology^1,3,4^.

Two specific deficits—increases in neural response variability and disruptions in oscillatory (periodic) activity—are linked to the onset and severity of deficits in schizophrenia^1^. For periodic signals, there is ample evidence that disruptions occur across multiple frequencies (for reviews see Newson & Thiagarajan, 2018^5^; Uhlhaas & Singer, 2010^1^). These patterns are typically explained as a series of band-specific changes in the theta (4-8 Hz), alpha (8-12 Hz), beta (12-30 Hz) and gamma (>30 Hz) bands. Disruptions in oscillatory bands may explain deficits in perceptual processing, and disorganized thought patterns^1^. However, the direction of these effects depends on the task itself and the specific sample^51^.

While changes to oscillatory patterns in patients with schizophrenia have proved robust in the experimental literature, there is scarce direct evidence regarding changes in non-oscillatory, response variability or “noise” in schizophrenia. This deficit exists because until recently separating aperiodic from periodic activity was difficult, or impossible. Recently, however, it has been shown that aperiodic activity can be reliably estimated from the EEG power spectrum^7,8^. Specifically aperiodic changes in the spectral domain are well modeled by a characteristic 1/*f^χ^* shape. That is, power in a spectrogram decreases as a function of frequency, where the exponent χ determines the steepness of the decline. This can be modified by altered excitation/inhibition balance in cortical layers, with increased inhibitory currents resulting in larger exponents and therefore steeper spectra^42^.

Traditionally, oscillatory power in a spectra was measured assuming the 1/f signal is constant. Emerging evidence, however, suggests aperiodic signals can vary in meaningful ways^8–12^. For example, healthy aging is associated with an increase in aperiodic activity, which was interpreted as “neural noise”^13^, and with changes in the overall shape of the power spectrum^8,14^. Furthermore, recent studies have reported alterations in aperiodic neural activity in mental illness. In patients with ADHD, spectral steepness was reduced on a dual performance attention “Stopping Task ‘‘ compared to healthy controls^15^. The shape of the power spectrum has also been found to be steeper in resting state EEG in patients with schizophrenia compared to controls^16^. This study highlighted the potential utility of using aperiodic signals to predict cognitive deficits in schizophrenia. When considered together with the results from Ostlund^15^, there may be an important difference in aperiodic activity that underlies cognitive tasks in schizophrenia.

The aperiodic activity perspective is particularly intriguing from an information theoretic perspective, which suggests that increasing aperiodic activity fundamentally decreases the efficiency of communications^17^, which is consistent with decreases in neural signal-to-noise in schizophrenia^18^. Given that the EEG is driven largely by the integrated postsynaptic currents converging onto a cortical region, and that aperiodic activity, in part, reflects the relative contributions of excitatory and inhibitory drive^42^, then significant shifts toward inhibition should fundamentally alter the communication efficiency of the cortical region while also manifesting as altered aperiodic activity, with steeper spectra resulting from increased inhibitory signals.

Modeling studies, however, suggest that aperiodic activity and oscillatory coupling are often mechanistically interdependent. For example, increasing noise can destabilize neural oscillations^19^. Likewise, variations in oscillatory power and phase can modulate background excitability^20^ and therefore alter neural communication.

Contrary to their independent mechanisms, periodic and aperiodic processes manifest as measurable signals which can confound each other in practice. That is, a change in aperiodic signal can confound attempts at measuring periodic power, unless care is taken to isolate each. With the development of *specparam*, overall spectral shape and band power may be measured separately. These independent measures convey information about different neural processes: the oscillatory or more traditional band measures, plus non-oscillatory, aperiodic background signal.

The current study investigates oscillatory and aperiodic activity differences between patients with schizophrenia and healthy controls during a target detection task. We first hypothesized that *measurements* of band-limited oscillatory power may be confounded by changes in aperiodic activity (H1)^22^. To separate these two factors we decomposed the power spectrum into aperiodic and periodic (band power) factors^7^. Thus, based on prior reports linking schizophrenia symptomatology to altered neural noise^21,55^, we hypothesized that patients with schizophrenia should exhibit steeper power spectra when compared to healthy controls (H2). We then undertook multivariate classification analysis of the factored EEG data to identify whether aperiodic activity or adjusted peak power was the most reliable predictor of schizophrenic clinical status in our data.

In our classification analysis we hypothesized two compatible outcomes. *1*) Band power would predict clinical status (H3); this result is consistent with the existing large literature on oscillations. *We also predicted that aperiodic power could predict status* (H4). We report that we find support for H2, H3, and H4. But note that contrary to our naive expectations, it was aperiodic signals that seemed the stronger and more consistent predictor of clinical status.

## Methods

### Participants

EEG and behavioral data were acquired for 60 (1 left-handed) participants recruited from two groups: patients with schizophrenia (SZ, n = 24, 7 female, age range 19-38) or control participants (n = 36, 16 female, age range: 19-41 years). Data originally acquired for 65 participants, but 5 from the clinical group were removed due to insufficient data, too noisy data, missing information. Patients with schizophrenia were referred by their treating clinician or were recruited from the community alongside the control participants. Patients met the DSM-IV criteria for schizophrenia-spectrum illness using the SCID interview^23^. Participants were compensated for their participation. To reduce long term effects of medication, only recent-onset patients were studied, with onset occurring within the last five years (see *Table 1*.). An exclusive inclusion of 5 years allowed a period for patients to adapt to their diagnosis, medication and/or treatment. Yet this period was not so long as to enter a window of long-term psychosis and medication use. Patients with schizophrenia were stable and continued treatment plans throughout the course of the study. All participants provided informed consent consistent with the University of North Carolina at Chapel Hill Institutional Review Board prior to their participation.

**Table 1.** Participant demographics and medication.

|  |  | **SZ** | **C** |
| --- | --- | --- | --- |
|  | *Age* | 24.71041667 | 27.74705882 |
| Sex | *Number Male* | 19 | 19 |
| Race | *White* | 19 | 22 |
|  | *African American* | 3 | 9 |
|  | *LatinX/HispanicX* | 0 | 2 |
|  | *Other* | 2 | 2 |
| Medication | *Antipsychotics* | 70.83% | - |
|  | *Anticonvulsants* | 22.73% | - |
|  | *Mood stabilizer* | 18.18% | - |
|  | *Lithium* | 13.64% | - |
|  | *Anticholinergic* | 9.10% | - |

### Experimental protocol

Data was recorded from a standard 10-20 electrode montage 32-channel EEG cap (Electo-Cap International) using NeuroScan 4.3.1 software and a SynAmps amplifier. Continuous EEG data were sampled at 500 Hz with a 0.15-70 Hz band-pass filter and 60 Hz notch filter. Electrode AFz served as ground and the right mastoid as reference. Vertical and horizontal bipolar electrooculogram (EOG) electrodes were placed at the outer canthi of each eye and above and below the right eye. Electrode sites were exfoliated and contacts were affixed with conductive gel. Electrode impedance was kept below 5 kΩ. Recordings were acquired in a sound-dampened chamber with participants seated 80 cm from the stimulus presentation monitor.

### Task

EEG was recorded while participants completed a target detection task (see Figure 1). Stimuli consisted of a letter (E, H, L, or P) or number (3, 7, 8, or 9) presented in either red or green typeface on a black background. Each participant was instructed to identify a target stimulus of the correct alphanumeric class and presented in the correct color. Letters were considered as targets, numbers as non-targets. Stimuli consisted of four combinations: targets were letter stimuli of the indicated color, stimulus mismatch non-targets were numeric stimuli of the indicated color, color mismatch non-targets were letter stimuli of the non-indicated color, and full mismatch stimuli were numeric stimuli of the non-indicated color. All bins occurred with equal frequency (25%). Stimuli were presented for 200 ms with a jittered interstimulus interval of 1500 ± 200 ms, during which a white fixation cross was presented. Participants performed 5 blocks of 100 stimuli each for a total of 500 trials. Overall, 125 trials of each condition were presented in a pseudorandom order. Participants were instructed to respond to the target stimuli with the left mouse button (index finger) and to all other stimuli with the right mouse button (middle finger). Early analyses showed that the three non-target stimulus types exhibited similar behavioral and physiological responses. Therefore, data from these stimulus conditions was combined by drawing equal numbers of trials at random from each of the three conditions such that the total count matched the number of target trials for each participant.

**Figure 1.**
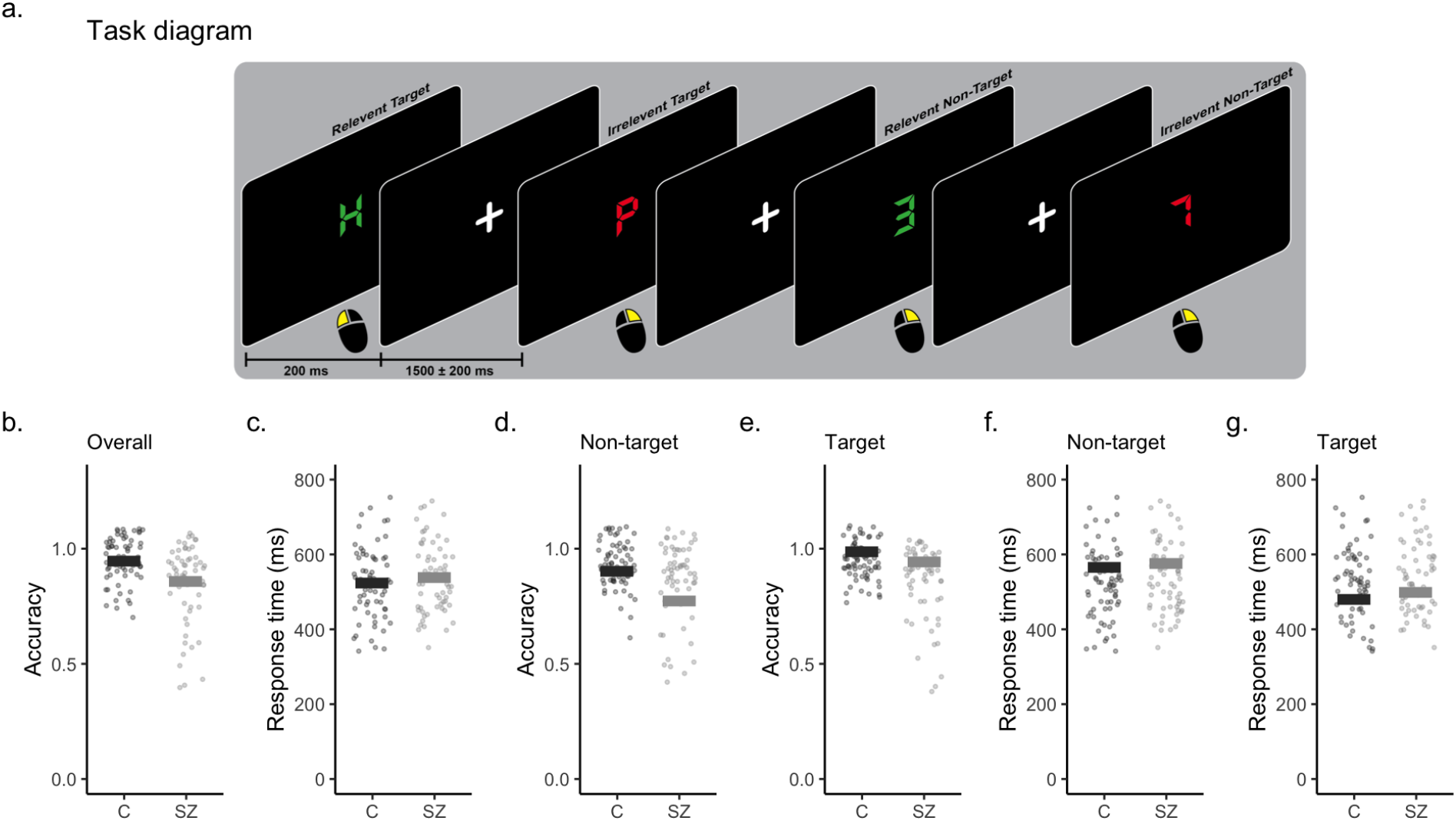
Task and behavior results. **a**) Task diagram. **b-c)**Accuracy and response time between the groups (control [C] and schizophrenia [SZ]). Comparison of mean (**d-e**) accuracy and **f-g**) response time between the two groups as a function of task conditions - non-targets versus targets.

### Data analysis

Continuous EEG data were imported to Matlab (v 2018b) and EEGLAB^24^ for processing. Data were high pass filtered at 1 Hz, provided channel locations, and submitted to automatic subspace reconstruction to identify and correct or remove artifactual data and channels (parameters were set to 8 s, 0.85 correlation, 20 SDs)^25^. Channels that were removed were then interpolated using the spherical method in EEGLAB (mean channels interpolated = 2.80). Data were segmented into epochs from −3s to 3s surrounding stimulus presentation to allow for frequency analysis. Epochs containing artifacts (as determined by a ±100 μV threshold, improbable, or high kurtosis data [+/-5 SDs]) were removed from further analyses. All subsequent analyses were performed independently for each EEG channel using only the last 3 seconds of epoch data, corresponding to periods of stimulus onset and the intertrial interval. During preliminary exploratory data analysis we considered all 30 channels (as in Figure 2–3). However, concerns about model overfitting--where the sample number approaches the feature number--prevented us from using all 30 channels in constructing our classifier. Instead we selected three channels for use during classification analysis (Fz, Cz, and Pz). We selected these three channels as prior experiments on target detection suggest this task relies on frontal and posterior neural generators and event-related activity in these areas is typically well-identified by simply monitoring frontal midline, central and posterior channels^48,49,50^. That is, Fz, Cz, and Pz.

**Figure 2.**
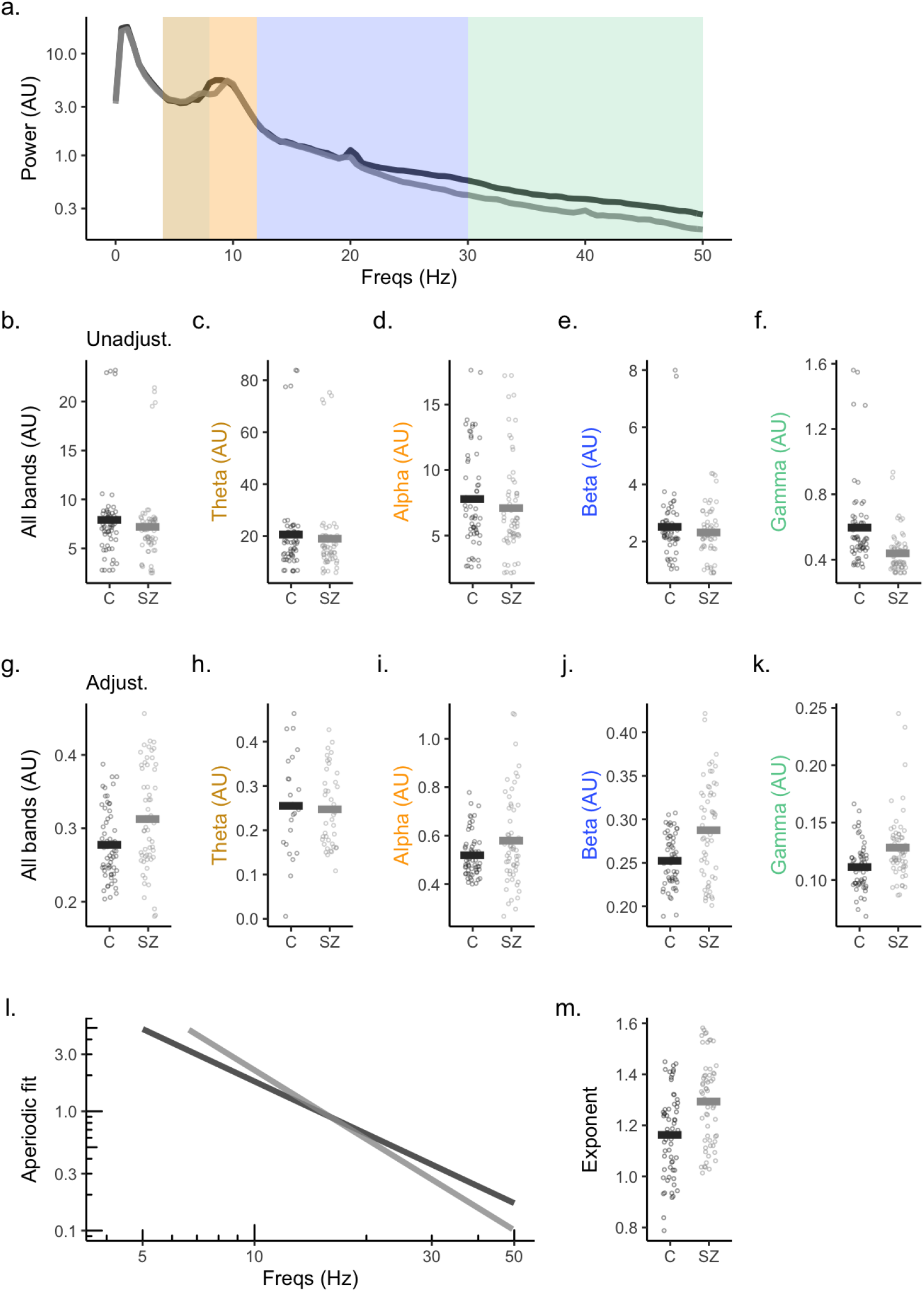
Oscillatory power and aperiodic activity in patients with schizophrenia (SZ; gray) and controls (C; black). **a**) Power spectrum (log transformed). Color coded regions represent oscillatory bands - theta (yellow, 4-8 Hz), alpha (orange, 8-12 Hz), beta (blue, 12-30 Hz), gamma (green, 30-50 Hz). **b-f**) Max band power across all bands. and for each band. Note: these values are the maximum power for each band *without* adjustment using the SpecParam algorithm. **g-k**) Max power for each peak detected by the SpecParam algorithm, *adjusted* for aperiodic contributions (see **l**), then binned into frequency bands. It is these power values which are used for all future analysis. **l**) Average aperiodic activity, with periodic activity removed. **m**) Exponent (steepness) differences between groups. In all plots dots represent individual values. Bars are the sample mean.

**Figure 3.**
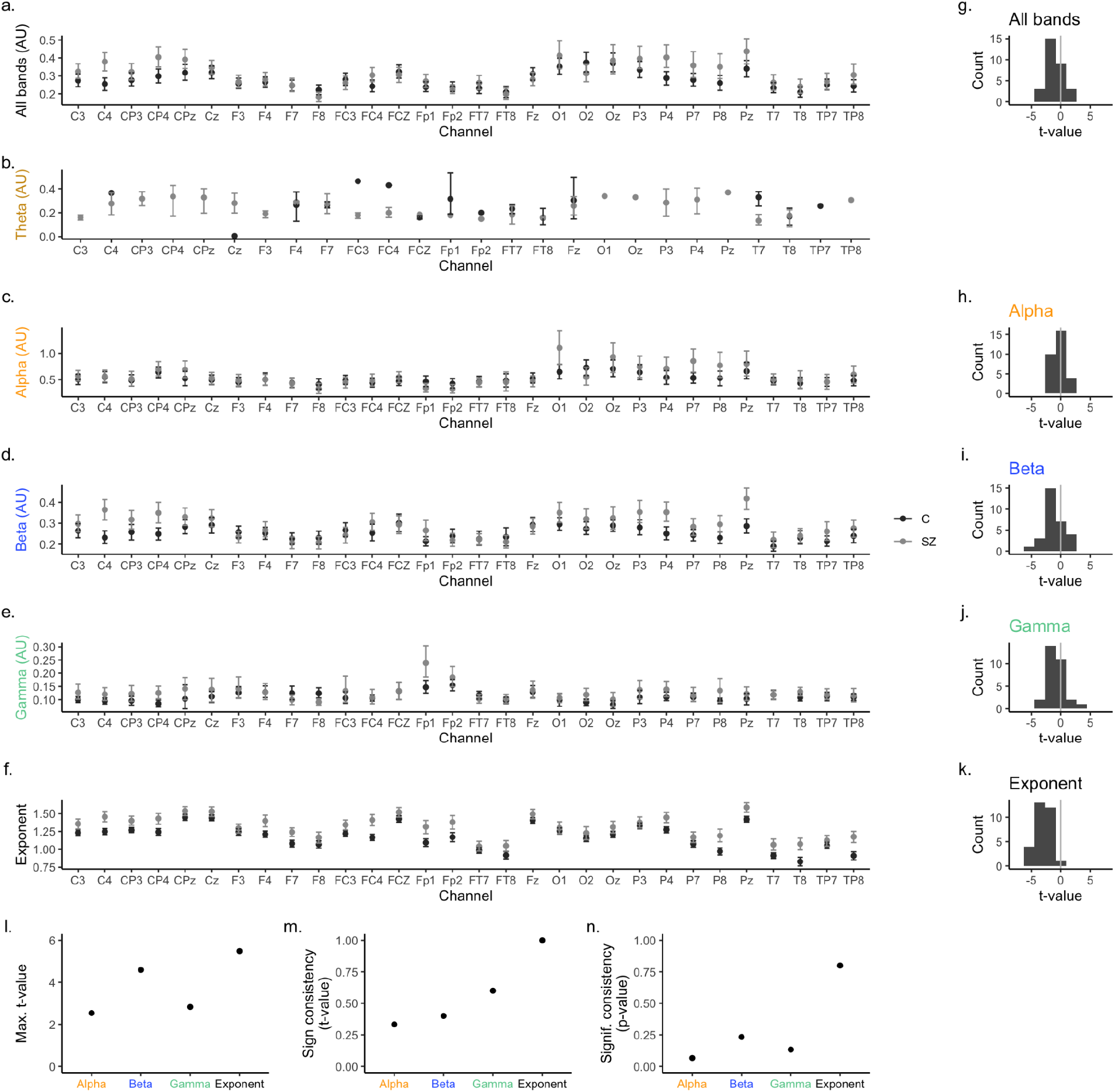
Electrode-level consistency and effect size. **a-e**) Band power for each of the 30 electrodes. Patients with schizophrenia (‘SZ’, gray) and controls (‘C’, black). **f**) Average spectral exponents for each of the 30 electrodes. Same conditions as above. **g-k**) Distributions of t-values for all electrodes (two-sided t-tests, between C and SZ conditions). The theta band had an insufficient number of peaks preventing reliable statistical testing (n=11, all participants). **l**) Maximum t-value for all electrodes. **m**) Consistency in the “sign” of t-values for all electrodes. **n**) Consistency of achieving significance for all electrodes, regardless of the direction of “sign” or direction of the statistical effect. Data in this figure was drawn only from the training dataset. See *Methods* for details on all metrics shown here, and for details on construction of the training dataset. In the histograms for panels **g**-**k**, the t-values shown were drawn from all channels and subjects.

Frequency domain analyses were performed using Welch’s method^26^. For each participant and channel, the PSDs of all epochs of each condition were averaged together. The aperiodic activity in each spectrum was estimated using SpecParam^7^ across the 4-50 Hz range. In brief, this algorithm decomposes the spectrum into an aperiodic component and a set of oscillatory peaks. The peaks are modeled as Gaussian functions. Each Gaussian peak is then taken to represent a single oscillation in the power spectrum. The maximum power from each of these peaks, and therefore each detected oscillation, was the basis of our measurement of band power. That is, traditional spectral analysis was performed then by binning the maximum power of each detected oscillation into the following “canonical” bands: theta (4-8 Hz), alpha (8-12 Hz), beta (12-30 Hz), and gamma (30-50 Hz). Participant performance was quantified by computing accuracy and response times (RTs).

### Statistical analysis

All statistical analysis and visualization was done using the R programming language^27^. Analysis code, and raw data is available at *[available post-publication]*. Oscillatory power measures were log transformed prior to statistical testing due to the log-normal distribution of oscillatory power^53^. Statistical testing used the full datasets to maximize sensitivity. That is, both training and reporting data (described below). These statistical tests did help select which bands to use in classification but beyond that in no way contributed to classifier hyperparameter tuning, testing or training. Prior to calculating summary statistics all 5% quantiles were calculated for power values. To remove the very large values sometimes present in power data, the 95% quantile was used to remove the top 5% of the data from further statistical analysis at the trial level to filter out a few large outliers. Power values that were adjusted by subtracting out the aperiodic signal, and those that were unadjusted, were treated separately during this removal procedure.

### Classification

We used Random Forest Classification to predict patient status (SZ or C) in a target detection dataset. To ensure a rigorous test of model performance, prior to undertaking any analysis 30% of both neural and behavioral data was randomly selected as a holdout or reporting set (which are equivalent terms here). The remaining 70% was used as the training set as part of a 5-fold cross validation procedure to tune model parameters. Performance analysis of the reporting set occurred after model tuning was complete. All normalization and rescaling of features was done separately for training and reporting sets, ensuring their independence. To estimate the reliability of classifier accuracy, an estimate of null performance (i.e., shuffled label performance) was used. To estimate chance performance, we resampled (with replacement) the group labels (C or SZ) generating a null distribution (see grey distributions in Figure 4). These distributions consist of 10,000 independent samples.

**Figure 4.**
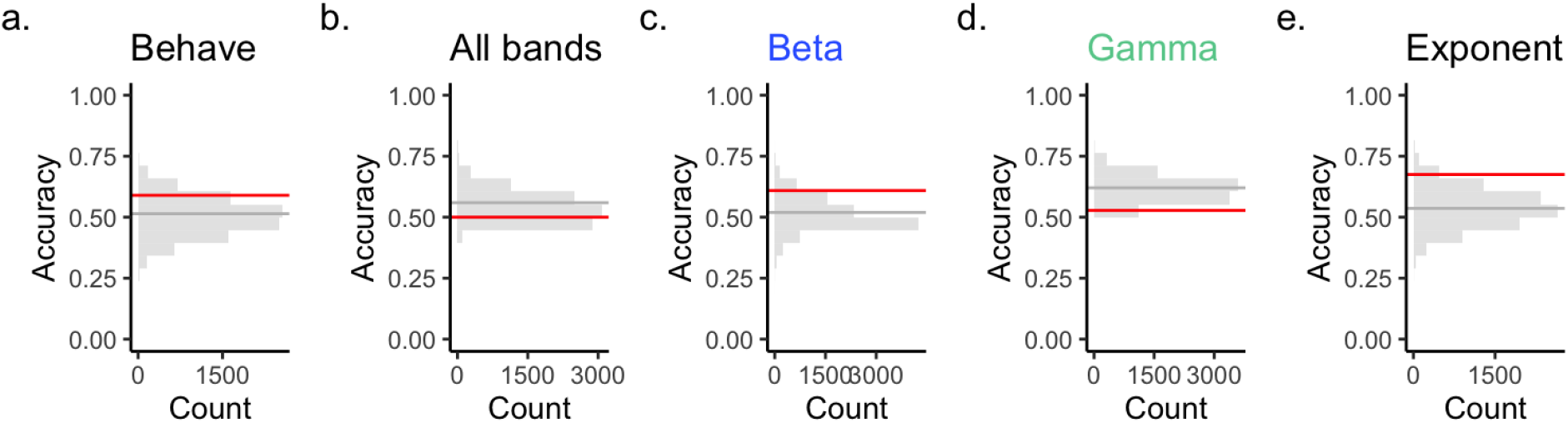
Classifier performance. **a**) Classification accuracy using only behavioral data **b**) Performance using all power from all bands **c-d**) Accuracy for two canonical oscillatory bands (beta, gamma, see Figure 3). **e**) Accuracy using aperiodic activity. Classification accuracy (red) was assessed on a hold out data set, comprising 30% of each data set. The performance above chance was estimated by comparing hold out performance to a label-shuffled null distribution (gray).

We focused only on the high frequency beta and gamma bands for classification analysis. The theta band was not included because our SpecParam algorithm detected too few theta peaks to build a meaningful classifier. Alpha was not included because little to no separation between C and SZ conditions was observed when carrying out our initial statistical analysis, see Figure 3**h**.

As noted above, we selected Random Forests^28^ for classifying both types of neural data - spectral power and aperiodic activity - and behavior. This algorithm is often performant and common in the neuroscience literature. Applying linear models instead to the neural data tended to reduce performance in the training set by 5-10% (not shown). Hyper-parameter choices are shown in Table 2. All preprocessing and machine learning analysis was done in Python 3.6 using the sci-kit learn package (v0.21.3)^29^.

**Table 2.** Tuned classifier hyperparameters.

| Features | Hyper-parameters |
| --- | --- |
| Behavior (accuracy, response time) | max_depth = 2<br>n_estimators = 5 |
| Aperiodic activity | max_depth = 2<br>n_estimators = 2 |
| Oscillatory power (all bands) | max_depth = 2<br>n_estimators = 2 |
| Oscillatory power (beta) | max_depth = 2<br>n_estimators = 3 |
| Oscillatory power (gamma) | max_depth = 2<br>n_estimators = 3 |

## Results

Our primary interest is to distinguish between patients with schizophrenia (SZ) and healthy controls (C), while both these groups undertook a consistent cognitive task. As this is the first of its kind study, we chose a simple target detection task. For two reasons detailed interactions between task-related effects (e.g. target versus non-target) and their relation to clinical status (C versus SZ) were deemed less important than simply separating C from SZ. First, in the long-term--beyond this one study--we seek a generally applicable biomarker. Second, building separate classifiers for complex task-status effects might lower the size of the training dataset to put the whole classifier at risk of overfitting. Nevertheless, we present some preliminary task-status interaction in the following section, along with the more critical analysis of overall clinical status.

### Task accuracy differs in patients with schizophrenia

Patients with schizophrenia had significantly lower accuracy (Mean 0.86 +/- 0.15 [SD]) compared to the control group (0.95 +/- 0.06 [SD]) (*t*(79.9) = 4.13, *p* = 8.9e-05; Figure 1**b-c**). Both groups had better performance for target trials compared to non-target trials (control (C): 0.99 +/- 0.02 [SD] versus 0.94 +/- 0.08 [SD]. *t*(47.15) = 7.26, *p* = 9.5e-09; schizophrenia (SZ): 0.94 +/- 0.08 [SD] versus 0.78 +/- 15 [SD]. *t*(47.15) = 5.19, *p* = 4.3e-06). Choice response time was not a significant indicator of group membership (t(126.93) = −0.87, *p* = 3.8e-01; 524.08 +/- 47.10 [SD] compared to 538.45 +/- 18.18 [SD]). Within groups reliable response time effects were observed, such that in control participants response times to targets were significantly faster (480.74 +/- 82.97 [SD]) compared those to the non-target stimuli (567.52 +/- 88.26 [SD]; *t(59.984) = −4.26, p = 6.7e-05*). Similarly, patients with schizophrenia were significantly faster in responding to targets (499.55 +\- 83.36 [SD]) than to non-target stimuli (577.5 +/- 85.53 [SD]); *t(59.984*) = −3.61, *p* = 6.3e-04; Figure 1**d-g**).

### Band-specific increases and decreases in oscillatory power

When peak power was assessed with the aperiodic component removed (“adjusted”), in the theta band there were no significant power differences between control and schizophrenia groups (*M*=0.26 +/- 0.12 [SD] versus *M* =0.23 +/- 0.096 [SD]; *t(42.6*) = −0.046, *p* = 9.6e-01). In the alpha range patients with schizophrenia also showed no significantly elevated power (0.57 +/- 0.37) compared to controls (0.52 +/- 31; *t(1172.4*) = −1.67, *p* = 9.4e-02). In the beta band, controls had significantly decreased mean power (*M* = 0.25 +/- 0.15 [SD]) than patients (*M* = 0.29 +/- 0.17 [SD]; *t*(3404.5) = −6.10, *p* =1.2e-09. As did gamma, where power was significantly increased in patients (0.12 +/- 0.07) versus controls (*M* = 0.11 +/- 0.06 [SD]; *t*(1085) = −2.51, *p* = 1.2e-02), See, Figures 2**h-k**. However, a mostly opposite pattern of results was observed in the unadjusted data. That is, power values extracted directly from the power spectrum without correcting for aperiodic changes or without explicit peak detection.

In unadjusted data the theta band showed no significant power differences between control and schizophrenia groups (M = 12.55 +/- 5.86 [SD] versus M = 12.54 +/- 5.51 [SD]; t(2668.1) = −2.8e-1, p = 0.7778). In the alpha range patients with schizophrenia also showed significantly decreased power (6.05 +/- 5.19) compared to control (5.91 +/- 5.68; t(3032.1) = −2.31, p = 2.1e-2). In the beta band, controls had significantly more power (M = 2.35 +/- 1.99 [SD]) than patients (M = 2.24 +/- 1.81 [SD]; t(2790.7) = 2.76, p = 5.8e-3. As did gamma, where power was significantly decreased in patients (M= 0.44 +/-0.52 [SD]) versus controls (M = 0.59 +/- 0.87 [SD]; t(2781.8) = 12.81, p = 2.2e-16), See, Figure 2**b-h**. Given the previously established independence of the aperiodic and periodic measures^7^ we use adjusted peak power from here on.

It has not escaped our notice that the different patterns of power seen between adjusted and unadjusted data, in the alpha, beta, and gamma bands, along with the presence of aperiodic effects (below), raises the possibility that other aspects of the existing EEG literature on schizophrenia might, in principle, show similar differences. It is also possible that this apparent “confounding” may help explain why the direction of oscillatory effects in schizophrenia appears task and context dependent^51^. That is, some studies may detect a mix of periodic and aperiodic effects, while others have unknowingly reported only aperiodic effects, while others reported pure periodic effects. These results underscore the importance of considering aperiodic activity in clinical samples.

### Aperiodic activity best predicts schizophrenia status

The power spectrum in patients with schizophrenia was significantly steeper (*M* =1.31 +/- 0.40 [SD]) compared to controls (*M* = 1.17 +/- 0.31 [SD]; *t* = −15.18, *p* = 6.0e-51; Figure 2**f-h**).

### Consistency and effect sizes across electrodes

To estimate the consistency of any differences in oscillatory power and aperiodic activity across electrodes, we visualized average measured values between electrodes. Visual inspection suggested that there was notable disagreement about the direction of effect in the theta, alpha, gamma, and beta bands. Some electrodes predicted band power should increase with schizophrenia status while other electrodes showed the opposite pattern--a band power decrease (Figure 3**a-e**). However, for all channels we observed that schizophrenia was associated with an increase in the steepness of the spectrum(Figure 3**f**). To approximate channel-level effect sizes, we calculated t-values for a comparison between C and SZ participants. These are shown as distributions in Figure 3**g-k**. Inspection of these distributions suggested that t-values for spectral exponents were both consistently larger than band power measures (compare Figure 3**k** to Figure 3**g-j**) and consistentlynegative. That is, aperiodic activity differences consistently had the same mathematical sign (Figure 3**f**).

We quantified these apparent differences in consistency in three ways. First, we measured the maximum t-value across electrodes for alpha, beta, gamma, and aperiodic measurements (Figure 3**l**). This serves as a proxy measurement for the strongest “local” effects we might expect to observe in some “best” subset of the electrodes. Here, the aperiodic measure gave the largest best effect by about 15%. Second, we tabulated the fraction of electrodes which had the same direction of effect or “sign” (Figure 3**m**). This “global” measure of effect direction indicated spectral steepness was perfectly consistent across all electrodes, having a value of 1. This corresponds to about a 100% increase compared to the next most globally consistent measure, which was gamma power. Third, we tabulated the fraction of electrodes which were significantly independent of the direction of effect (Figure 3**n**). Again, aperiodic activity was the most “globally” consistent by about 65%. Significance here was set using an uncorrected threshold of *p < 0.05* (Bonferroni correction did not change the relative pattern of results we observed).

Spectral power varies between cortical regions and can be used to identify individual subjects, and tasks^30^. We therefore employed a spatial classification approach, based on a Random Forest classifier, to predict clinical groups (i.e., control or schizophrenia) from either oscillatory band power or aperiodic activity (see *Methods*). We contrasted these outcomes with predictions made using behavioral data.

The classification model based on aperiodic activity (Figure 4**e**) achieved the highest median classification accuracy (0.68) compared to a model using all oscillatory band data, which was at chance levels (0.50) (Figure 4**b**). Aperiodic activity also outperformed beta (0.62) or gamma (0.53) power alone (Figure 4**d**). The model based on behavioral response (*i.e*.,response time and response accuracy) performed better than either of the oscillatory models (0.60; compare Figure 4**a** to **b-d**). Behavioral predictive performance was, however, inferior to aperiodic activity (0.60 compared to 0.68; Figure 4**a** to **e**).

## Discussion

We hypothesized that measurements of band power would be confounded by changes in aperiodic activity (H1), and that patients with schizophrenia would exhibit steeper power spectra consistent with increases in cortical inhibition (H2). We further hypothesized two outcomes during classification analysis. Band power would predict clinical status (H3) and/or aperiodic activity would predict status (H4). Our findings further suggest that aperiodic activity is a stronger predictor of schizophrenia. Furthermore, band power measures done without accounting for aperiodic activity appeared to be strongly confounded (confirming H1).

Aperiodic activity was also seen to be a more reliable predictor than measurements derived from oscillatory bands. Given both its strength of effect and its consistency between electrodes, we suggest large-scale study of aperiodic activity under a wide variety of cognitive tasks is warranted by our work. We believe aperiodic activity may be able to serve as a general-purpose biomarker for schizophrenia^16^’.

### Behavior as a diagnostic tool

Behavior is commonly used as a diagnostic factor in schizophrenia and patients show behavioral deficits on multiple domains (attention, memory, working memory, and so on). However, these deficits are not considered biomarkers in and of themselves because they lack the predictive and prognostic sensitivity. As such, behavior is impaired, but this impairment is not sufficiently consistent enough for use as a biomarker. A good biomarker for schizophrenia would align with tenets laid out by Weickert^52^ and the NIH which include (1) diagnostic to classify as having a disease, (2) prognostic to make predictions on who will develop a disease, or (3) theranostic to predict an individual response to a particular therapy. The limits of behavior-based diagnostic approaches motivate our use of EEG-based predictors. Likewise, for any EEG-based predictor to hope to succeed as a useful predictor it ought to perform better than behavioral analysis alone.

### Oscillatory power differences in context

The direction and significance in band-specific differences observed in the current report are broadly consistent with prior work (see Figure 2). The only exception is the lack of a group difference in theta power. However, all other bands displayed a significant separation based on clinical status.

There is ample evidence that gamma activity is disrupted in schizophrenia. Interpreting these disruptions, however, is difficult. Current results suggest gamma *oscillations* are highly stimulus, task, and context dependent^31–33^, though it is important to note that power in the gamma band may not always reflect the presence of a true oscillation in that band^34^. Even when present, the power of gamma oscillations fluctuates strongly in time, and is suppressed by task-irrelevant factors^31,32^. We interpret the fact that gamma alone was a poorer predictor of disease status than aperiodic activity to be due in part to this variability. We additionally note that there was a strong alpha peak present in the power spectra in our data (Figure 2**a**). Like gamma, this difference was significant, however the strength of alpha’s predictive capacity using spatial variations in power (a classification accuracy of 0.56 +/- 0.12) was still well below that of spectral shape (0.71 +/- 0.11) and also well below behavior-based predictions (0.65 +/- 0.12).

### Aperiodic activity and oscillatory power may be complementary

In this target detection dataset we observed that aperiodic activity better predicts schizophrenia compared to band-specific power changes. Combined with previous results in cognitive aging^8^, ADHD^15^, Autism Spectrum Disorder^35^, and schizophrenia^16^, we argue that non-oscillatory, aperiodic neural activity can serve as a useful biomarker in clinical populations. Furthermore, we report the first difference in schizophrenia during a cognitive task. The ample literature reporting band specific anomalies in patients with schizophrenia complement the range of studies that suggest oscillatory activity and coupling is relevant for healthy cognitive function. However, given the growing body of work in aperiodic activity, future work should consider isolating changes in band-specific power from changes in aperiodic activity.

Based on the strength of our classification results, we expect aperiodic activity to prove a reliable predictor of schizophrenia status outside this single experiment. While we do not expect aperiodic activity will explain the wealth of band-specific variations across the literature, we believe that measurements of band-limited power and aperiodic activity may prove to be complementary pieces of information that will improve the utility of electrophysiology measurements as clinical biomarkers. Indeed, both oscillations and aperiodic activity may be needed to understand and computationally model the complex biological interactions that generate schizophrenia pathophysiology. For example, an increase in gamma power that occurs in line with a chronic increase in inhibitory conductance (see below) may have an overall suppressive effect on neural excitability^36^ whereas in a more homeostatic condition, a gamma oscillation can instead act to increase neural gain^37^. That is, the functional role of gamma oscillations may change as inhibitory conductance increases. Oscillations with a strong inhibitory effect may act to gate or suppress information transmission ^36,38^, whereas gamma oscillations operating in a more balanced regime may instead amplify information flow^37^.

### Steeper spectra may reflect increased inhibitory conductance

The shape of the power spectrum is described by a 1/f power law distribution. We focus here on the range from ~4-50 Hz. In this range, exponent (χ) values vary from to 1-4^10,11^. The larger the χ the steeper the power spectrum will appear. Recent modeling results which link changes in power spectral exponents/steepness to changes in synaptic activity^39^ and excitatory and inhibitory currents^40^. These efforts, which relate synaptic fluctuations in the membrane voltage to local field potentials (LFP), are supported by recent combined recordings of LFP and single units in human subjects showing that slower inhibitory time constants dominate the LFP^41^.

In response to experimental results, we have developed a computational model to infer changes in excitatory-inhibitory balance directly from spectral shape in LFP recordings^42^. In this model, increasing the inhibitory conductance, or the time constant of inhibitory kinetics, increases steepness. Conversely, decreasing the proportion of inhibitory activity, or increasing excitatory conductance, decreases steepness. That is, our model suggests the increase in the spectral exponent we report here may be a direct consequence of increased inhibitory conductance. This increase is presumably a homeostatic response to the decreased number of inhibitory neurons associated with schizophrenia. Within a limit, increasing the strength of individual synapses can compensate for the overall reduction in inhibitory interneurons, however this may reach a pathological tipping point over time.

If the observed change in spectral shape is caused by a loss of inhibition and the corresponding homeostatic adjustment, this may explain why aperiodic activity is the superior predictor of disease status. Loss of inhibition appears as a widespread and chronic phenomenon in schizophrenia. That is, while excitatory-inhibitory imbalance ultimately affects oscillatory coupling, oscillatory changes depend on a number of external factors (*e.g*., attention, stimulus contrast, size, task context, goals). These external factors may act as dynamic “confounds” that are shared between control participants and patients with schizophrenia undertaking the same cognitive task. Meanwhile aperiodic activity may reflect a more fundamental physiological index - inhibitory conductance - allowing for the observed improvement in classification performance. Power laws can however also arise from a large number of physical sources^43^. In neural electrophysiology, aspects of the power spectral exponent have been attributed to shot and brownian noise^44,45^, and to network connectivity^9,10^. Others use the 1/*f*^χ^ pattern to suggest the brain is operating like a dynamical system tuned to a self-organized “critical point”^46,47^.

#### Aperiodic consistency

Our belief in the importance of consistency in the aperiodic signal is based out of pragmatic measurement concerns. It is often the case that the band power changes in disease states at rest, and especially during task performance, will show a high degree of variation at the subject level. This is true even though the average effect appears consistent between cohorts, as in a t-test. Population level consistency is adequate for laboratory studies where the aim is population inference. But if the aim of biomarker development is to identify and quantify individuals at large, then individual variation becomes extremely important. We therefore find it noteworthy and promising that the aperiodic signal is so consistent because it increases the likelihood that a classifier, like the one we use here, would succeed in practice in the clinic, and moreover, that it could have broad utility regardless of the EEG setup since it was observed across channels.

### Drug effects, and other important caveats

We selected patients with schizophrenia who were diagnosed within the last five years in an effort to minimize drug effects contaminating the electrophysiological results. However both the aperiodic and band specific changes we report here may nevertheless be due, perhaps in part, to pharmacological confounds. Controlling for this possibility in a chronic condition like schizophrenia is difficult. Ultimately we will need to confirm these results in drug free individuals and in animal models. However more elaborate experiments require an initial robust result, which is what we report here.

### Summary

We have reported results for a single target detection task. Despite our relatively large patient pool, these results require confirmation in other tasks and, ideally, in other laboratories. The current study adds to the growing literature which shows the importance of investigating aperiodic activity in clinical settings in general, and in schizophrenia specifically.

## Acknowledgements

Work supported by grants 5R01MH158251; 5-P50-MH164065

